# Self-supervised retinal thickness prediction enables deep learning from unlabeled data to boost classification of diabetic retinopathy

**DOI:** 10.1101/861757

**Authors:** Olle G. Holmberg, Niklas D. Köhler, Thiago Martins, Jakob Siedlecki, Tina Herold, Leonie Keidel, Ben Asani, Johannes Schiefelbein, Siegfried Priglinger, Karsten U. Kortuem, Fabian J. Theis

## Abstract

Access to large, annotated samples represents a considerable challenge for training accurate deep-learning models in medical imaging. While current leading-edge transfer learning from pre-trained models can help with cases lacking data, it limits design choices, and generally results in the use of unnecessarily large models. We propose a novel, self-supervised training scheme for obtaining high-quality, pre-trained networks from unlabeled, cross-modal medical imaging data, which will allow for creating accurate and efficient models. We demonstrate this by accurately predicting optical coherence tomography (OCT)-based retinal thickness measurements from simple infrared (IR) fundus images. Subsequently, learned representations outperformed advanced classifiers on a separate diabetic retinopathy classification task in a scenario of scarce training data. Our cross-modal, three-staged scheme effectively replaced 26,343 diabetic retinopathy annotations with 1,009 semantic segmentations on OCT and reached the same classification accuracy using only 25% of fundus images, without any drawbacks, since OCT is not required for predictions. We expect this concept will also apply to other multimodal clinical data-imaging, health records, and genomics data, and be applicable to corresponding sample-starved learning problems.

## Main

Ophthalmology is a field that is pioneering artificial intelligence applications in medicine and has recently experienced many promising results that can significantly change the future of eye care. The field has benefited from recent advances in deep learning,^1,2^ particularly in the case of deep convolutional neural networks (CNNs) when applied to large data sets, such as two-dimensional (2D) fundus photography, a low-key imaging technology that captures the back of the eye. These images can be taken using a smartphone and are available in a standardized fashion, often in very large quantities.^3^ Using data from diabetes screening programs and biobanks, cardiovascular risk factors, presence of diabetic retinopathy, and even gender, can be predicted with a high degree of accuracy.^4,5,6^ More recently, CNNs were applied to three-dimensional optical coherence tomography (OCT) of the retina to segment tissue layers and predict retinal referral decisions.^7,8^ In all these approaches, large data sets and ever-developing leading edge models from natural image domains have been used, often requiring cumbersome annotations as well as substantial computational complexity. Regarding the widespread clinical use of deep learning, which often needs to be embedded in mobile and on-device applications,^9^ the issue of large-scale samples with expensive annotations, as well as computationally expensive models, must be addressed.

While machine learning concepts, such as transfer learning and domain adaptation, have made significant progress in terms of enabling the use of deep-learning algorithms on smaller data sets,^10,11^ features learned from natural images, such as those in ImageNet,^12^ do not necessarily transfer meaningfully to the medical domain, where visual features, resolution, and output targets may differ considerably.^9^ In addition, ImageNet-based model architectures are specifically designed for predicting 1,000 output classes, and are often heavily overparameterized in the context of medical imaging problems.^9^ Natural image features, combined with non-optimal model architectures, inhibit transfer learning from ImageNet for medical image data sets. To address these problems, we believe effective transfer learning, using pre-trained medical data models, is required, along with greater model flexibility. However, a large, annotated medical image data set, comparable to ImageNet dimensions, does not currently exist.

The concept of self-supervised learning (SSL) may offer a solution to this problem. The notion of SSL is to determine useful representations from unlabeled data by solving pretext tasks. A pretext task is an inference problem for which labels can automatically be generated. Through training according to these labels, the model learns to extract relevant (visual) features.^13^ Examples of such an approach include restoring color to grayscale images, predicting correct rotation, or recovering the correct semantic ordering of natural images.^13,14^ Recent SSL approaches using more abstract formulations by, for example, learning context-consistent, patch-wise image representations, indicate that it is possible to match fully supervised ImageNet pre-training performance in effective transfer learning tasks using natural images.^15^ Drawing inspiration from recent work on SSL to recover partially masked input signals through the use of multimodal observations,^16,17^ we implemented large-scale cross-modality SSL in the medical domain, while keeping the necessary annotations effort low.

For the successful application of SSL it is necessary that the chosen pretext task has to learn embeddings which are meaningful for the respective downstream tasks. Defining an effective pretext task for medical imaging data is particularly challenging since relevant pathology-related features are often represented through subtle and small-scale phenomena. These render conventional SSL tasks less effective since they are tailored toward the presence of dominant objects in natural images. This also holds true in the field of ophthalmology, where subtle changes in the eye can refer to significant differences. For example, in clinical routine examinations, patients often undergo unnecessary treatment for neovascular age-related macular degeneration (nAMD), using anti-vascular endothelial growth factor (anti-VEGF) medication.^18^ The underlying disease is actually macular telangiectasia, a rare retinal disease that resembles nAMD OCT in appearance, but does not respond to this specific treatment.^19^ Other subtle and rare retinal changes include retinal angiomatous proliferation or polypoidal choroidal vasculopathy,^20,21^ which is often mistaken for nAMD, thereby preventing the correct treatment. A successful SSL pretext task must ignore an image’s large, non-informative aspects and focus on representing disease encoding subtleties.

In this paper, we propose a novel SSL pretext task for medical data in ophthalmology, by encoding shared information between two entirely different high-dimensional medical modalities, namely, OCT and infrared (IR) fundus photography. More specifically, we first extracted fundus retinal thickness maps from co-registered OCT (Fig. 1a) with little labeling, then predicted these maps from the IR fundus images directly, using a deep CNN-based model (DeepRT) (Fig. 1b). We showed that this SSL pretext task learned a fundus representation containing disease-relevant features, thereby enabling transfer learning, significantly improving downstream disease classification using limited samples. We demonstrated this novel deep-learning pipeline (Fig. 1) on a large data set, provided by the Munich University Eye Hospital,^22^ and showed that the OCT-derived retinal thickness map was predicted accurately directly from fundus photography. To evaluate this pretext task, we ensured that the DeepRT learned disease-relevant signals by numeric evaluation and tested our predictions against the ground truth, via a consortium of doctors, within a typical disease screening setting. We then employed the learned feature representations for transfer learning onto a public diabetic retinopathy detection data set (Fig. 1c), and showed that the self-supervised network outperformed both random and ImageNet initializations for diabetic retinopathy classification in a scarce training data scenario, while reducing model size by more than two orders of magnitude.

**Figure 1:**
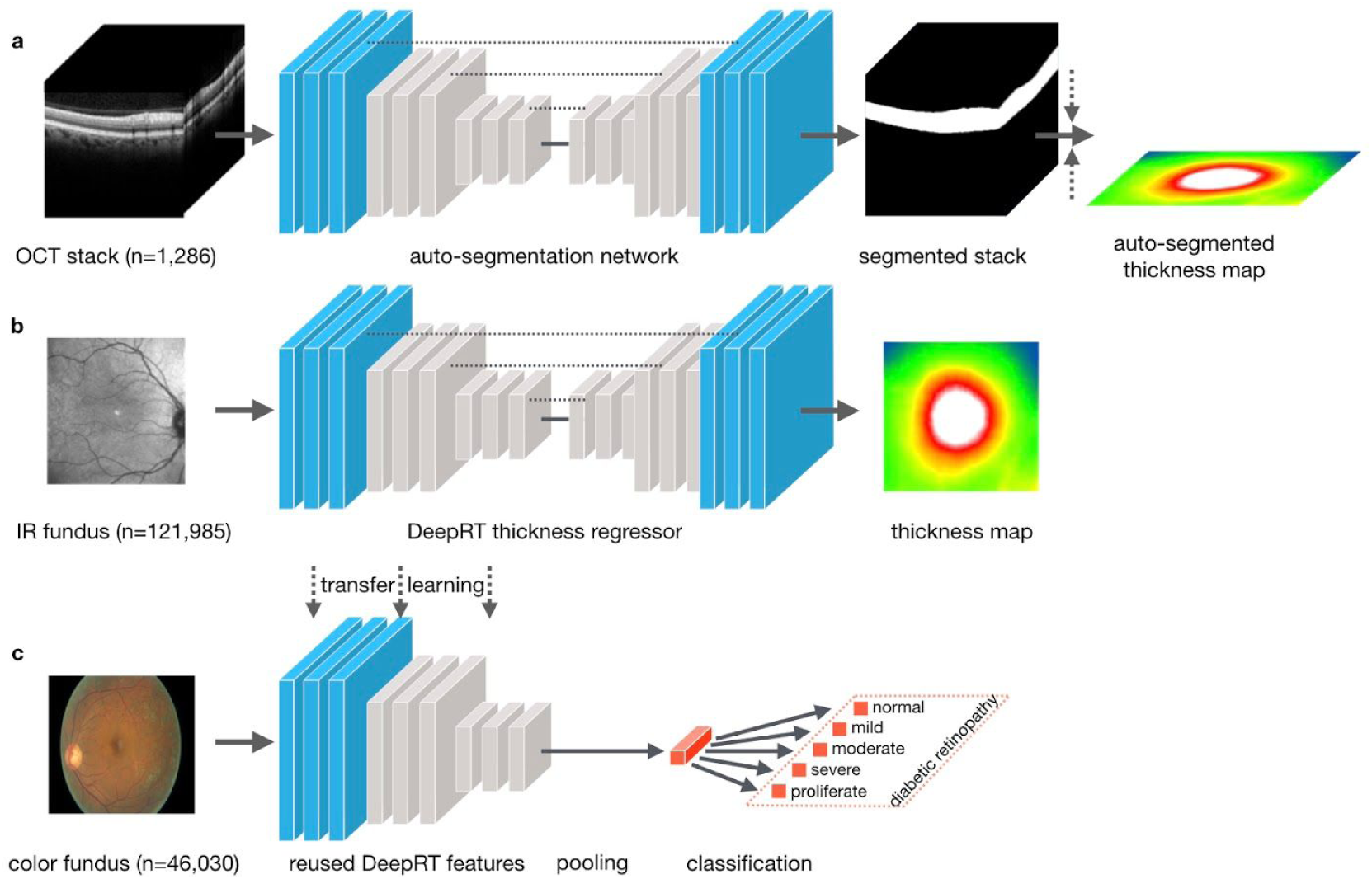
Cross-modality self-supervision workflow: absorbing physical devices and algorithms into a single neural network learning disease, using relevant features from unlabeled data. **a.** Deep learning powered OCT segmentation enabled ground truth retinal thickness maps. **b.** Ground truth thickness maps used for cross-modal prediction of retinal thickness directly from fundus images. **c.** Features learned in b were transferred to an independent, single modality downstream classification task.

## Results

### Deep learning allowed for gaining robustly segmented OCT retinal tissue

To prepare training data for the thickness predictor (DeepRT), we set up a U-net based^23^ deep-learning system for automatic pixel-wise tissue segmentation (Fig. 1a). The algorithm was trained and validated on 1,009, and tested on 277, manually annotated OCT brightness (B)-scans across two different devices, Spectralis^24^ (device I) and Topcon 3D^25^ (device II). The algorithm performed according to a high mean intersection over union (IoU)^26^ of 0.94 on the training, validation, and test set alike (Fig. 2a). Examples of segmentations from devices I and II show how the algorithm was able to segment OCTs from different devices with high accuracy (Fig. 2a, b). The algorithm was then applied to segment the total set of 121,985 volume scans, each with a stack of 49 OCT images, from 20,995 unique patients.

**Figure 2:**
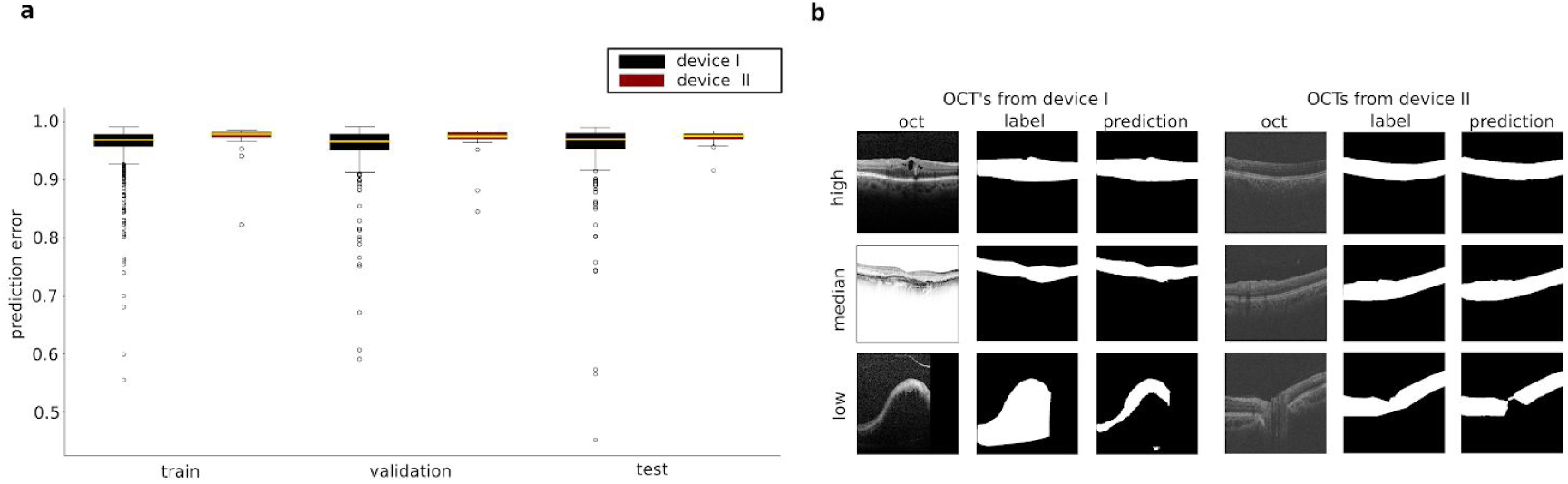
Accurate device-agnostic OCT thickness segmentation. **a.** IOU scores on training, validation, and test set for device I and II. **b.** Randomly selected OCTs from device I with high (0.98 IoU), low (0.45 IoU), and median (0.97 IoU) performance, and from device II with high (0.98 IoU), low (0.91 IoU), and median (0.97 IoU) performance.

### DeepRT accurately predicted high-resolution thickness maps directly from fundus images

Once all OCT scans had been segmented, we set up an SSL pretext task using DeepRT to predict thickness directly from the fundus photographs. To evaluate the DeepRT, we calculated the mean absolute error (MAE) and average deviance percentage on an independent test subset of 17,969 scans. The deviance percentage was calculated as the relative difference, with respect to the ground truth value, for each pixel. The MAE and deviance percentage were presented for all test records, test records with an observed ground truth thickness above 400 μm in the central subfield (CSF) region, or overall thickness map, and for test records, including the known presence of edema or atrophy (Fig. 3a). DeepRT’s predicted retinal thickness on average with a 33 μm deviation from the ground truth (Fig. 3a), which is on average less than 10% deviance with respect to each individual pixel. Thickness maps observing measurements of 400 μm or thicker in the overall or CSF region had MAEs of 50 ± 27 μm and 61 ± 32 μm, respectively. This corresponded to a 13% and 15% deviance. In addition to pixel-wise validation, we compared the average values over the clinically relevant score that is common in ophthalmology, namely, the nine macular sectors, based on areas defined in the Early Treatment Diabetic Retinopathy Study (ETDRS)^27^ (Fig. 3b). Finally, we captured thickness information with low percentage deviance, between 5%–10%, and high spatial awareness in cases of normal, as well as pathological, fundus images of diabetic macular edema (DME), vein occlusion and atrophy (Fig. 3c, d).

**Figure 3:**
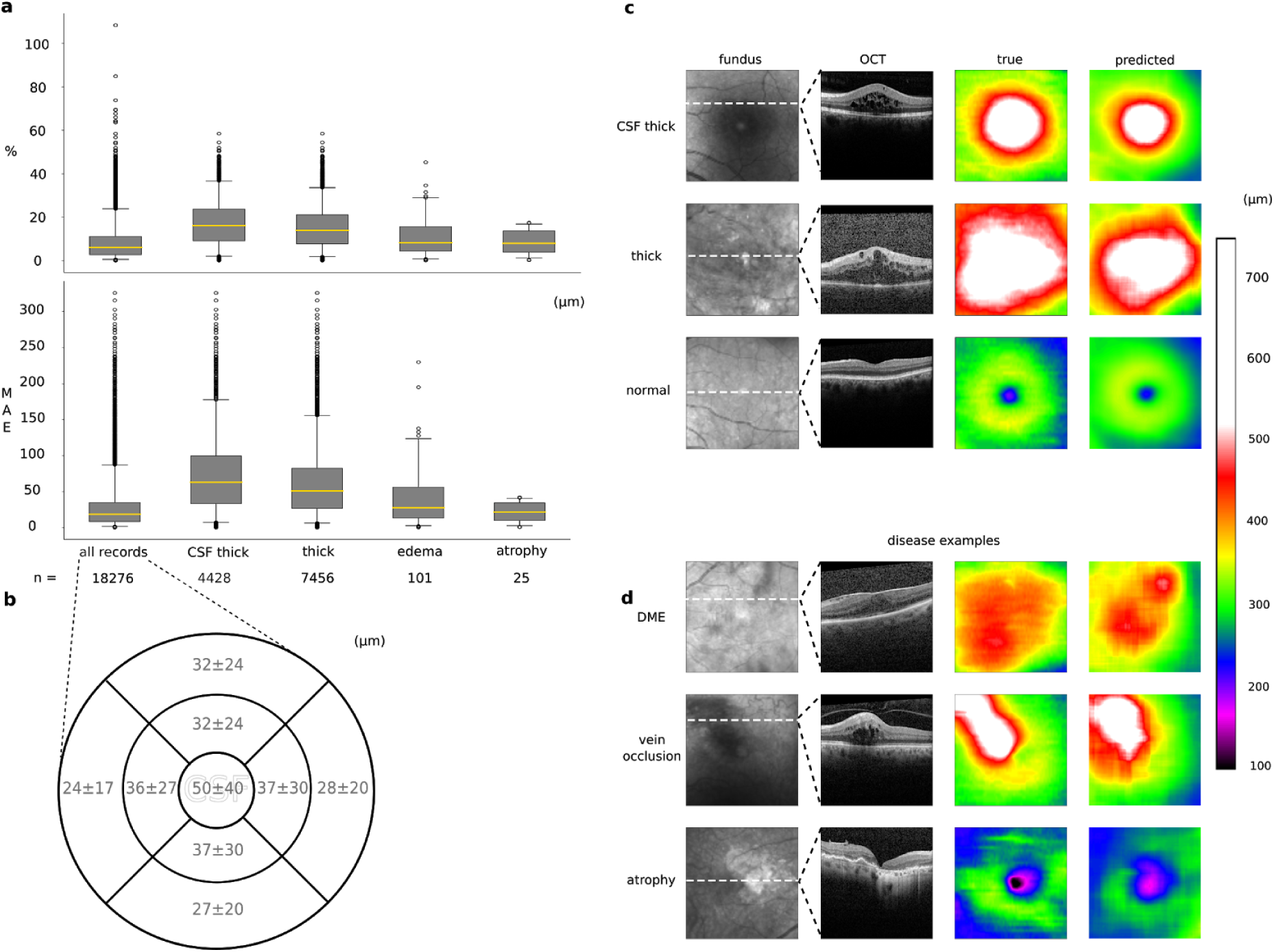
DeepRT consistently predicted retinal thickness across devices based on fundus images. **a.** Relative error and MAE plots for all, CSF thick, overall thick, edema, and atrophy test records. **b.** Spatially resolved errors across the ETDRS grid. **c.** Examples from the test set showing accurate thickness predictions in relation to OCT segmented thickness maps d. Examples of particular disease profiles found in real world clinical data.

### Clinical decision supported by predicted thickness

For additional evaluation of whether DeepRT learned disease-relevant representations, in addition to the standalone utility of the predicted thickness maps, we presented 261 randomly selected examples, actual and predicted, and employing various thickness profiles, to three retinal specialists (J.S., T.H., and T.M.), who were tasked with classifying the presence of macular thickening (Fig. 4a). These specialists were randomly shown real and predicted thickness maps, individually for each patient, and the information that two maps belonged to the same patient was not disclosed. On 79% ± 6% of the maps, the specialists all came to the same conclusion for both true and predicted maps, with regard to detecting thickening, and on 69% ± 2% for non-thickened retinas (Fig. 4b). For examples of true and predicted thickness maps causing different outcomes see supplementary information. Inter-doctor alignment rate was 89% ± 6% in the case of macular thickening; that is, on average, in 10% of cases, the predicted thickness maps caused the specialists to reach different conclusions than from the OCT-based ground truth. Finally, we asked the specialists to evaluate 261 patient records for detectable edema and atrophy using only the fundus, or the fundus and predicted thickness map available. The association was then modeled using a linear mixed effect model, accounting for each specialist as a covariate (see methods section). The ground truth diagnosis, the so-called *gold standard*, was determined using full OCT scans alongside fundus images, as well as patient history available at the eye clinic, and noted in electronic medical records. Macular thickening was defined if the central retinal thickness (CRT) in OCT was more than 400 μm, and atrophy if the CRT was lower than 250 μm. The resulting per class receiver operating characteristic (ROC) curve showed a higher area under the curve (AUC), detecting atrophy when the specialists had access to the predicted thickness map, which was not the case for detecting edema, where the AUCs were roughly the same (Fig. 4c).

**Figure 4:**
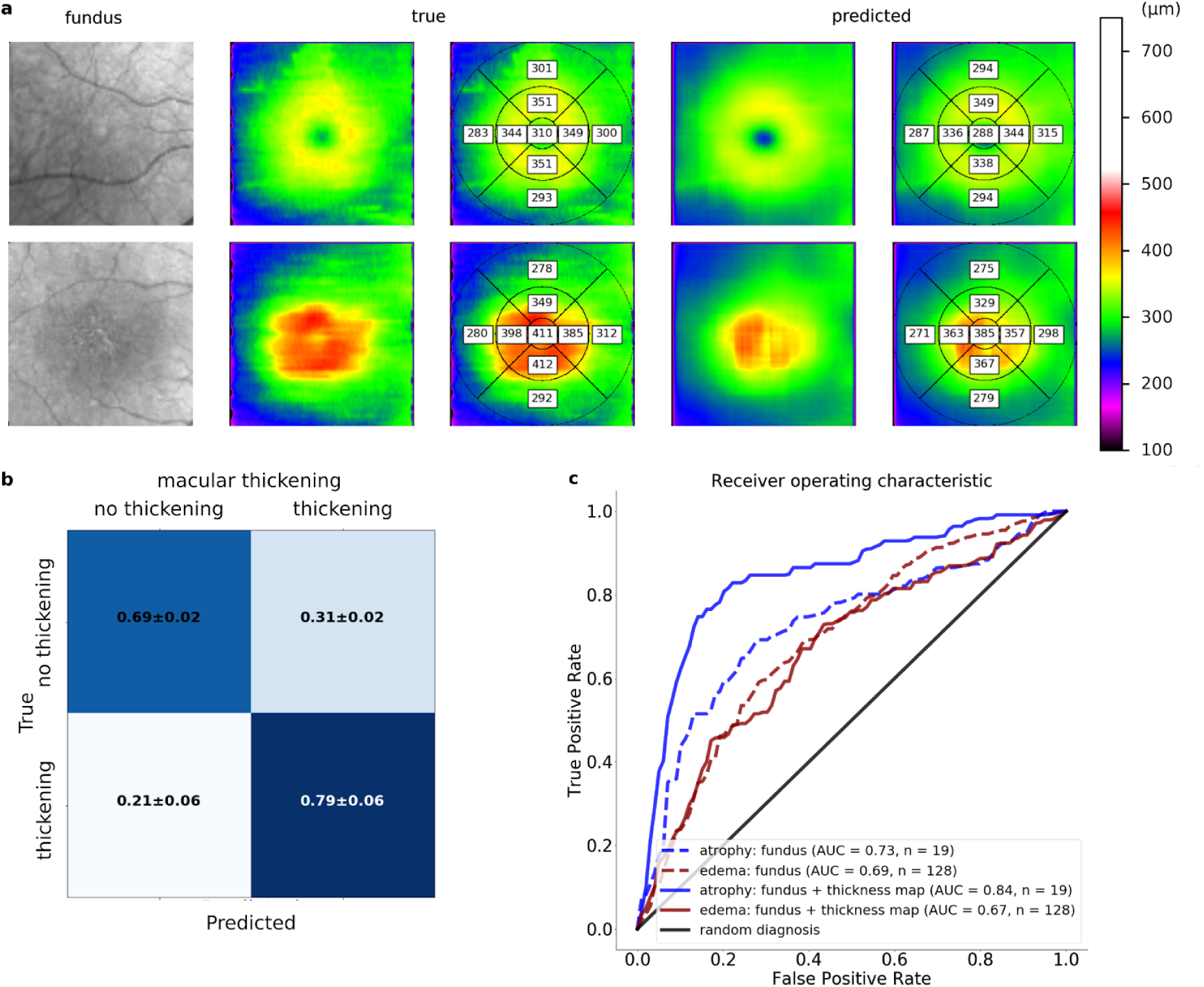
Inferring the gold standard from answers with or without access to a predicted thickness map, demonstrating that the proposed method increased the diagnostic capability of physicians. **a.** Example of true and predicted thickness maps given to specialists for determining the presence of macular thickening, edema, or atrophy. **b.** Alignment rates between true and predicted thickness maps when detecting macular thickening. **c.** ROC-curve for the mixed model, inferring the gold standard from answers, with or without access to a predicted thickness map.

### Self-supervised learning improved training and test performance in the context of classifying diabetic retinopathy

To address our original hypothesis, i.e., that SSL may lower the need for annotated samples in ophthalmology, we studied the effect of transferring weights learned in the DeepRT encoder to classify diabetic retinopathy on fundus images in an independent cohort of patients. The DeepRT thickness prediction allows flexible network design choices and is herein constructed as a ~ 125,000 parameter model, which is extremely lightweight compared to the full ResNet50 of ~23 million parameters. As the heavily parametrized ImageNet models, optimized for the natural images domain, render transfer learning onto medical data sets inherently inhibitive, this flexibility is an important feature of the DeepRT SSL pretext task and enables optimal medical task model design. To evaluate the DeepRT initialization, we repeated training five times with random and DeepRT initializations of the small CNN architecture against a state of the art Imagenet pretrained Resnet50 (CNN), using a Kaggle diabetic retinopathy data set provided by EyePACS,^28^ a free platform for retinopathy screening. We used the initial 35,124 training images and an independent public test data set of 10,906 images. Furthermore, to evaluate the more realistic and lighter sample setting, the above procedure was repeated on five data partitions, each excluding parts of the data, thus reducing the data set size to 25%, 10%, 5%, and 3% of the original set, stratified by class. For evaluation, we used weighted accuracy as a primary metric. Furthermore, weighted binary precision and recall, cohens quadratic kappa as well as cross entropy was observed (see supplementary information). Following extensive hyper parameter comparison, we found that the DeepRT-initialized model outperformed other initializations, particularly on the smaller partitions, against both the random lightweight and the ResNet50 ImageNet initialized models (Fig. 5a). Additionally, given its ideal lightweight architecture, the DeepRT initialization only needed 8,781 labeled examples (25%) to reach the same weighted accuracy, compared to when the equivalent model was randomly initialized, effectively replacing 26,343 annotated color fundus photographs. Across all partitions, the DeepRT initialization required substantially fewer annotated examples than the standard random initialization (Fig. 5b).

**Figure 5:**
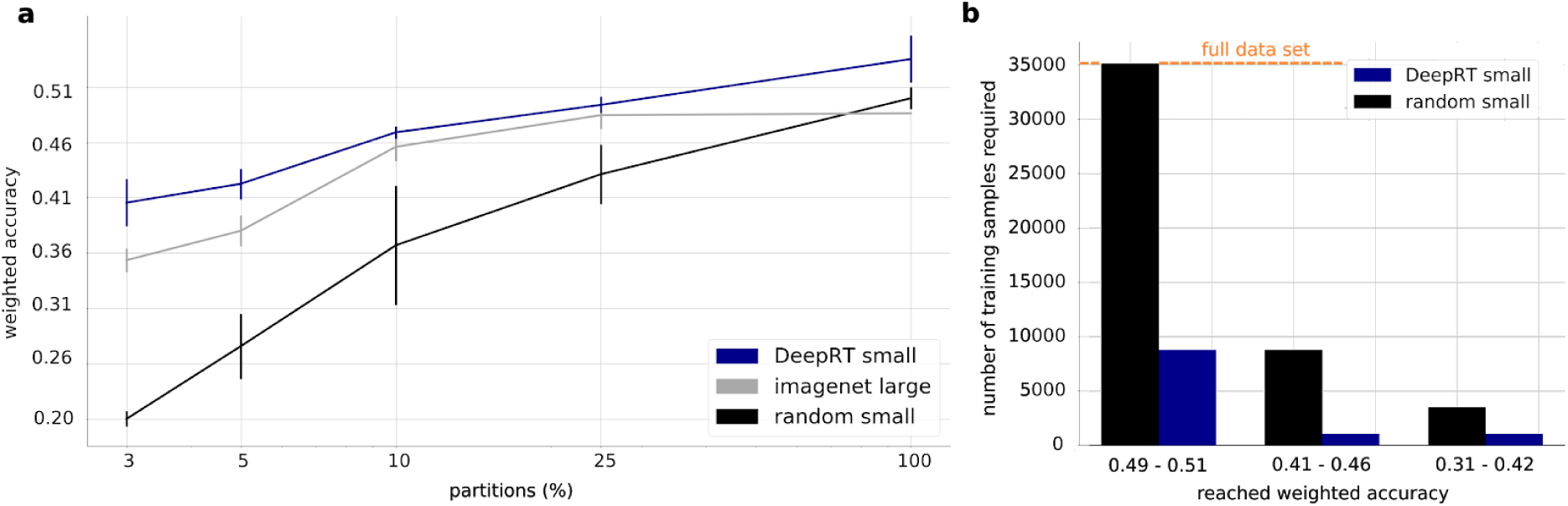
Self-supervised learning reduced annotated sample need four-fold. **a.** Average weighted accuracy and error bars displaying inter-model standard deviation across five models, trained on each partition, displayed on x axis in logarithmic scale, for random, ImageNet, and DeepRT initialization. b. Required number of annotated examples for achieved weighted accuracies of a random classifier as read from panel a, and corresponding number of samples needed for the DeepRT initialization to achieve the same result.

## Discussion

We showed how the presence of multimodal information, namely OCT and fundus IR images, can be exploited to reduce labeling efforts greatly, in a bid to achieve high accuracy on the downstream task of diabetic retinopathy classification. In our cross-modal approach, we used the fact that different medical signals can be learned from different modalities with varying difficulty. In the case of OCT, retinal thickness was directly extractable by training an efficient, leading-edge, U-net-based architecture to segment the retinal layer. The 1,009 annotations necessary for model training were generated in less than one week of annotation effort for a single clinician, after which the model reached a high mIoU of 0.94, allowing for the complete automated set-up of the efficient DeepRT SSL pretext task.

Since the SSL’s effectiveness depends on DeepRT’s capability to learn disease-relevant features, multiple evaluations of the predicted thickness maps were made. While small numerical differences (Fig. 3a) serve as a good indication of successful learning, they do not directly indicate to what degree DeepRT predictions can substitute for OCT-calculated thickness maps, arguably the most important goal, or test whether DeepRT has learned the variations in the fundus that encode the pathology-relevant thickness information from the OCT. To do this, we proceeded to evaluate our predictions in a typical screening setting to determine whether trained clinicians observed the same pathologies in predicted and ground truth thickness maps (Fig. 4). Our specialists achieved a high (79% ± 6%) alignment rate between predictions and ground truth maps when detecting macular thickening, particularly considering the inter-doctor variance of 89% ± 6%. Together with numerical evaluation (Fig. 3), the DeepRT proved its ability to capture disease-relevant features in most cases (Fig. 4b). Furthermore, when analyzing the standalone utility of our thickness maps, providing our panelists with the fundus only or fundus and predicted thickness map, we observed a considerable improvement in detecting atrophy in the retina (Fig. 4c). Depending on the cause, atrophy of the fundus is often non-trivial to observe. This is particularly true in generally reduced nerve fiber layer atrophy, as observed in neural degeneration or glaucoma. DeepRT thickness maps can help identify patients with these diseases since they can represent both pathologically thick and thin examples from fundus images. Surprisingly, specialists were able to detect edema, or lack thereof, from fundus photographs only; adding a thickness map did not make a difference in this regard. Thickness maps would be less important in long-standing edema, which create characteristic changes in the retina and therefore could be visible in the fundus. However, subtle edemas are difficult to identify in fundus imaging only. Our results indicates that a large part of retinal thickness information is encoded in both modalities (OCT and IR fundus) (Fig. 4c)

In the final stage of this work (Fig. 1c), we showed that learning this shared, cross-modal information in a self-supervised manner is an effective approach for initializing a diabetic retinopathy classification model using color fundus images. Remarkably, our self-supervised network steadily outperformed random and ImageNet initializations by a significant margin, despite the fact that the model was derived from training on small-field-of-view infrared images (Fig. 5a). With only 25% of the training data, the DeepRT-initialized algorithm nearly recovered the full weighted accuracy reached, compared to training conducted on the complete data set, reducing labeling efforts by a factor of four. Specifically, we showed that the proposed three-stage process can replace 26,343 expert diabetic retinopathy annotations (Fig. 5b) over an equivalent, randomly initialized model, with 1,009 relatively simplistic semantic OCT annotations, which medical staff trained at a lower level can achieve accurately. In addition, the process also reached an even higher classification accuracy than leading-edge transfer learning from ImageNet, while allowing a 200-fold decrease in model size, which is critical for the mobile and on-device applications that are often necessary for deploying deep-learning models in real scenarios. These results are supported by work in the research community, which indicates that, in medical imaging, features of interest are often subtle, represented within small areas, and therefore difficult to discover. Furthermore, model architectures suitable for medical imaging problems appeared to be considerably different from those used on ImageNet, making general features and models learned from natural images less useful for increasing medical data efficiency.^9^

Our model training strategy includes two other benefits over most multimodal approaches,^29,30^ which are typically structured around absorbing multiple modalities in a single model, in order to improve classification accuracy. First, although our training strategy relied heavily on multimodal data sources, it used only OCT information for pre-training purposes. The final model operated completely independently of OCT data at inference time; that is, we did not require an OCT measurement as input for the final diabetic retinopathy prediction model, which operates on a single color fundus image. This presents the benefit that our training strategy does not present additional complexities for clinical application (e.g., data fusion, missing data, and increased runtime), which are associated with multimodal modeling approaches. Second, we were able to learn from disjointed data sources, in the sense that we were able to effectively use patient OCT information to improve color fundus models on a completely different data set, with no patient overlap. We expect that this indirect use of multimodal patient information in the form of self-supervision will be particularly useful for deep learning in the medical domain, where multiple measurement modalities, coupled with disjointed patient groups, render simultaneous inclusion in shared models impossible and, as a result, exclude important studies and data sets.

### Study limitations and future outlook

Cross-modal, self-supervised pre-training incorporates medical knowledge and intuition in the form of the regression of retinal thickness, which is known to reflect a large portion of the relevant pathology for diabetic retinopathy. While incorporating this medical prior is one of the method’s core strengths, it also limits the methods general transferability to similar medical challenges with limited existing knowledge since the procedure requires extensive collaboration between medical staff and software engineers to conceptualize, implement, and evaluate a suitable pretext task. Furthermore, we only evaluated a single pretext task in this work. Cross-modal self-supervision between the fundus and OCT modality offers several possible extensions. For example, to additionally predict other OCT quantifiable clinical features such as epiretinal membrane detachments, directly from fundus photographs, could improve the effectiveness of capturing disease-relevant features. Further, new methods such as Cycle-Consistent Adversarial Networks^31^, have shown great success in image-to-image translation and could be applied to OCT and fundus pairs, potentially enabling even more effective self-supervised learning. Finally, recent progress on guided model architecture design has enabled more efficient models to be trained on natural images.^32^ These models were not evaluated in this work nor in previous studies on transfer learning onto medical images.9 Guided model design could offer additional efficiency gains from both Imagenet and medical SSL pre-training, enabling smaller and more accurate models on medical data.

## Online methods

In the next section, we explain the primary method-related information for this project. First, a brief explanation of the OCT and fundus image modality in ophthalmic practice is presented. Then, we explain the data sets, followed by the implemented algorithms.

### OCT and fundus images

Optical coherence tomography is a three-dimensional (3D) volumetric imaging technique that measures the reflection of infrared light in human tissue at a spatial resolution of less than 5 μm.^33^ A typical OCT examination yields an infrared image of the patient’s fundus and an accompanying co-registered stack of OCT images, providing a 3D view of the patient’s retinal morphology. This tomographic information is used to compute retinal thickness maps, which provide retinal experts and ophthalmologists with important information about pathologies and abnormalities in the eyes of their patients. Variations in this imaging modality are key to distinguishing and classifying various forms of macular diseases.^34–37^ Due to the high-resolution tomographic view provided by OCT, the fundus and OCT pair are routinely obtained at specialized eye clinics today; therefore, this information exists in large volumes.

### Tissue segmentation data set

The tissue segmentation data set comprised 1,286 OCT B-scans; 1,066 of the scans were obtained from the LMU eye clinic’s standard Spectralis OCT device, where each scan was selected from a different patient and annotated by a team of four doctors using an open source software called LabelMe (v3.16.1).^38^ Additionally, 110 publicly available OCT images were obtained from the Duke Enterprise Data Unified Content Explorer^39^ and from Golabbakhsh et al. (2013),^40^ respectively, all with provided annotations. The Duke repository OCTs were also obtained using standard Spectralis OCT, while for Golabbakhsh et al the Topcon 3D OCT-1000 device was used. The images were randomly split into 744 training, 265 validation, and 277 test images, with no patient overlap.

### Thickness prediction data set

The LMU eye clinic data set consisted of fundus and OCT pairs of 121,985 eyes, from 20,995 unique patients, generated from patient visits between 2012–09–24 and 2018–12–04. After filtering out faulty and too-low-quality records (see supplementary data), the data set consisted of 96,824 OCT volumes from 19,812 unique patients. These were segmented, superimposed, and linearly interpolated for thickness map calculation (see supplementary data). The DeepRT was then trained and evaluated using the 96,824 filtered fundus and thickness map pairs.

### Screening evaluation data set

The screening evaluation data set comprised OCT scans of 261 different patients, which were randomly selected according to the following criteria: only one eye from each patient could be included. Additionally, OCT scans of all diagnoses were included. Half of all scans showed no pathological changes, whereas the other half showed pathologies. Scans with thickening were overrepresented since this feature is more common. The data set was then analyzed by one clinician (K.U.K.) for correct alignment, and the correct category was decided on (normal, thickened, or atrophic).

### Kaggle diabetic retinopathy data set

The public diabetic retinopathy data set used for transfer learning (Fig. 1c) was taken from a previous Kaggle competition.^28^ The data set consisted of 35,126 color fundus images for training, and 10,906 images from the public test set for evaluation. The images were classified into five different stages of diabetic retinopathy: 0: none; 1: mild; 2: moderate; 3: severe; 4: proliferative diabetic retinopathy.^41^

### Tissue segmentation algorithm

Where tissue segmentation was concerned, a U-net architecture neural network was used, as in Ronneberger et al. (2015),^23^ with batch normalization and rectified linear unit activations after each convolutional layer, and added drop-out after each max pooling layer. As thickness information is easily extractable from the OCT modality, standard training configurations and preprocessing were used (see supplementary technical notes).

### DeepRT algorithm

The DeepRT network was modeled as a deep pixel-wise regression network. It consisted of an encoder and a decoder (Fig. 1b), the former a lightweight ~ 125,000 parameter model consisting of six units, each with two residual blocks containing skip connections, created with transfer learning for medical image tasks in mind. The decoder was a regular decoder, similar to a structure using coupled de-convolutional processes and sets of convolutional operations to process and increase data to input size (see supplementary technical notes).

### Linear mixed effect model

The responses from specialists detecting edema, atrophy, and/or normal patient records (Fig. 4c) were modeled using a linear mixed effect model. This is a logistic regression model in which the specialist is modeled as a covariate. Thus, each answer and specialist were considered covariates, and the outcome was gold standard values. This was subsequently modeled for all records where the gold standard was determined as edema, atrophy, or normal, as well as for fundus and fundus with predicted thickness map, individually.

### Kaggle diabetic retinopathy detection (DRD) transfer learning

When transferring the DeepRT weights, and benchmarking against random and ImageNet initializations, DeepRT and randomly initialized models were lightweight (~125,000 parameter) compared to ImageNet ResNet50 (~23 million parameter) models. During training, images were preprocessed using Gaussian blurring (sigma parameter 10), circle cropping, channel wise mean subtraction, and intensity standard deviance scaling, calculated from the training data. Images were augmented by random flipping and rotation, hyperparameter optimization of learning rates, learning rate decay schedules, and individual momentum for each configuration. Training was stopped manually inspecting loss convergence.

## Supporting information

complete supplementary text and figures.

## Acknowledgements

We thank the administrative teams at Helmholtz Zentrum Munich and LMU’s University Hospital (LMU-UH) for the fast data-sharing agreement. LMU-UH data protection officer G. Meyer for constructive and fast approval of data processing. We acknowledge the previous work of M. Müller for setting-up and maintaining the data warehouse at LMU-UH, A. Anschütz for its continuation. R. Wolff and A. Babenko for creating the electronic medical records in our hospital and C. Kern for the medical oversight. M. Rohm and I. Manakov for fruitful discussions on our data.

## Author contributions

O.G.H. developed the deep learning models and the data analysis pipeline. F.J.T. and N.K conceived the study. K.U.K. lead the data acquisition and data interpretation. T.M, J.S, T.H., L.K., B.A., J.S. did image annotations and screening evaluations. N.K supervised the study with F.J.T and K.U.K. F.J.T., N.K., O.G.H., and K.U.K. wrote the paper. O.G.H., N.K., K.U.K. and F.J.T. contributed to the interpretation of the results. All authors read and approved the final manuscript.

## Competing interests

F.J.T. acknowledges support by the BMBF (grant# 01IS18036A and grant# 01IS18053A), by the German Research Foundation (DFG) within the Collaborative Research Centre 1243, Subproject A17, by the Helmholtz Association (Incubator grant sparse2big, grant # ZT-I-0007) and by the Chan Zuckerberg Initiative DAF (advised fund of Silicon Valley Community Foundation, 182835). F.J.T. reports receiving consulting fees from Roche Diagnostics GmbH and Cellarity Inc., and ownership interest in Cellarity, Inc. and Dermagnostix.

N.D.K reports ownership interest in Hellsicht GmbH.

K.U.K declares no competing interests.

O.G.H declares no competing interests.

