## Supplementary material for "Self-supervised retinal thickness prediction enables deep learning from unlabeled data to boost classification of diabetic retinopathy": complete supplementary text and figures.

### Supplement

#### Supplementary note 0: Calculation of high resolution thickness maps

Below proceeds an explanation of the thickness map calculation. The process consists of thickness **vector calculation**, **superpositioning** and **interpolation**.

#### Vector calculations

First, all 49 OCT images of an eye were segmented. The segmentation were then used to calculate OCT specific 1D thickness vectors. One thickness vector was calculated by subtracting the distance in pixels between the top and bottom pixel classified as part of the retinal tissue along every position on the x - axis in a segmented OCT. If a patch of the OCT image was unsuccessfully segmented, causing parts of the thickness vector to show zero values, the patch is assigned the mean thickness values from the closest two thickness measurements in the thickness vector. If the the segmentation fails at the farmost right or left of the B-scan, then the values of the most left or right segmentation is used for imputation. Thereby ensuring that as few parts as possible of the thickness grid is assigned zero values yielding vastly incorrect measurements for parts of fundus image thickness maps.

#### Superpositioning

Positional and scale arguments were contained in the eye clinic export for every OCT. Exact start and end positions on x and y axis positions with respect to the fundus image were used to superimpose every OCT onto a fundus sized grid. Then available scale arguments were used to convert the pixel measured thickness values into µm, a more natural measure for clinicians.

#### Interpolation

Once all thickness vectors were superimposed and scaled to µm the values in between each thickness vector could be linearly interpolated, forming a high resolution thickness map on the area on the fundus where OCT images were obtained.

#### Supplementary note 1: Data exclusion

Due to the large number of data points generated from a real world practice the quality of the data varied substantially and automated filtering methods were required as manual revision was to cumbersome.

#### Data format and incompleteness filtering

The first filtering criteria applied was to use OCT stacks containing 49 B-scans only obtained from the macular region. Then, filtering was done by data incompleteness. Data incompleteness refers to missing or faulty positional and scale arguments in the image export.

If an OCT stack did not contain information of where each B-scan was obtained on the fundus, the calculation of the thickness map was not possible.

### Data quality filtering

In order to filter out records where OCT scans had too low quality average cross axis total variation of the calculated thickness maps was used. Cross axis total variation is simply the average total variation calculated across both the x and y axis. As the difference in thickness values in a correctly segmented thickness map should not vary too much, an efficient filtering threshold was found.

Filtering was an important step in curating the data as real world datasets typically contains noise and quality issues. While the goal of filtering as many bad records as possible was optimized, the balance between filtering out faulty records while retaining as much of the good data as one can was crucial. The threshold 4.19 of cross axis variation was found by visual inspection of randomly selected filtered and not filtered thickness maps. A clear separation between "hacky"  and clearly incorrect thickness maps was found for images above the threshold of 4.19 while still retaining ~ 100,000 examples (Fig. 6).

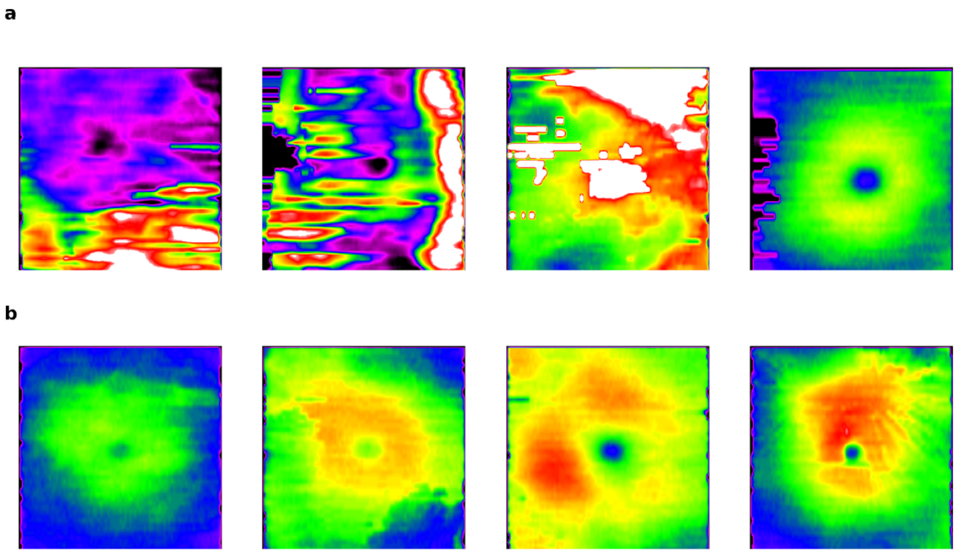


**Figure 6: Randomly selected examples of filtered and not filtered thickness maps:** ***a.*** *Filtered thickness maps with larger then 4.19 µm in cross axis total variation.* ***b.*** *Kept thickness maps with less than 4.19 µm in cross axis total variation.*

### Supplementary note 2: Technical notes on thickness segmentation algorithm

### Preprocessing and training

For training the U-net tissue segmentation algorithm the Adam optimizer was consistently used and initialized with the default values for parameters beta 1 and beta 2 with 0.9 and

0.999 respectively, for further details see.[1] The network parameters were initialized using the He normal initialization[2] with default values and the network was optimized using the dice loss[3]. During training, each OCT was resized to 256x256 pixels using bilinear interpolation, scaled between 0 and 1, rotated and flipped and trained with tuned hyper parameters. After a grid search for typical values the optimal set of parameters were found to be dropout: 0.2, learning rate: 0.001.

**Data availability:**
All annotated OCT images for tissue segmentation from the LMU eye clinic will be made available in the future.

**Supplementary note 3: Technical notes on DeepRT thickness prediction**

**Preprocessing**
The fundus images and thickness maps where resized from 768x768 to 256x256 pixels to make training faster. Further, the thickness maps were only obtained from a center square part of the fundus, the remaining margins where cropped out. Gaussian blurring was applied to every fundus image with sigma parameter set to 10. Finally each fundus image was normalized to values between 0 and 1 and thickness maps were also normalized by a factor of 1/700, making almost all values between 0 and 1.

**DeepRT architecture**
The DeepRT architecture used was a combination of a light weight encoder and a U-net style decoder with the same architectural specification as in the original U-net paper. The encoder consisted of 6 residual units, each with 2 residual blocks accumulating a total of 42 convolutional layers and a total 126,373 parameters. As in the U-net model, there are feature layer skip connections between the corresponding convolutional layers after each unit in the encoder and deconvolution operation in the decoder. As in the tissue map segmentation network, batch normalization and drop out was added in the decoder after each convolutional layer. With a heavily parametrized decoder, the full DeepRT model consisted of 32,411,985 parameters.

**Training details**
The model was trained for 30 epochs, until convergence on the validation set was observed. A learning rate schedule was set to decrease the learning rate an order of magnitude from 0.001 to 0.0001 after 20 epochs. Stochastic gradient descent was used for optimization with added momentum set to 0.99. The loss function was mean absolute error (mae) which was also used as the model selection criteria on the validation set. Training took 6 hours to complete and was stopped manually inspecting loss convergence.

**Implementations**

All the implementations were done using python and tensorflow keras and are available here: https://github.com/theislab/LODE/tree/master/DeepRT. The models where trained on a GeForce GTX TITAN X GPU from Nvidia.

**Supplementary note 4: Examples of true and predicted thickness maps causing different outcomes in detecting thickened and non thickened retinas:** Testing the predicted thickness maps in a screening like setting (Fig. 4) showed the potential utility of predicted as a substitute to OCT derived maps. Surprisingly, especially detecting non thickened retinas yielded only 69 % alignment rate between predicted and OCT maps. Observing randomly selected cases where such misalignment occured, (Fig. 7) shows that in ambiguous examples, where slight thickening might occur, the maps can be difficult to evaluate. In Fig 7b, DeepRT clearly misses a small edema, most likely causing the predicted map to be differently evaluated.

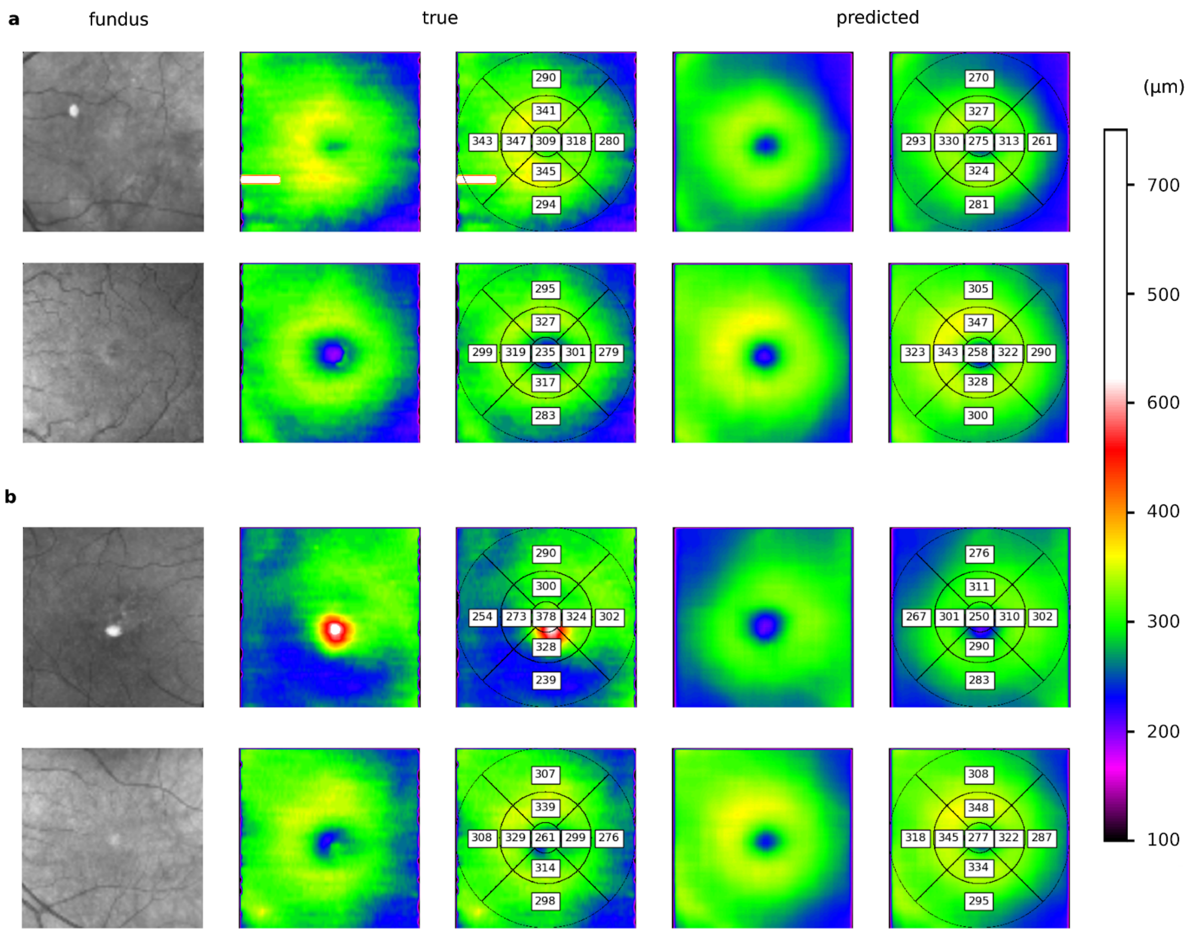


**Figure 7: True and predicted thickness maps yielding different screening outcomes:** *Each row is the fundus, true and predicted thickness map for one patient a. Randomly selected examples where the true, OCT derived, thickness map was determined as non thickened and the predicted as thickened by at least on specialist. b. Randomly selected*

*examples where the true thickness map yielded thickened and predicted thickness map non thickened screening results by at least one specialist.*

**Supplementary note 5: Further analysis of transfer learning onto diabetic retinopathy images**

Binary weighted recall show an even greater difference in performance between the DeepRT initializations, random and state of the art imagenet. DeepRT consistently outperforms the other intilizations across recall, precision and the common cohens quadratic kappa (Fig. 8).

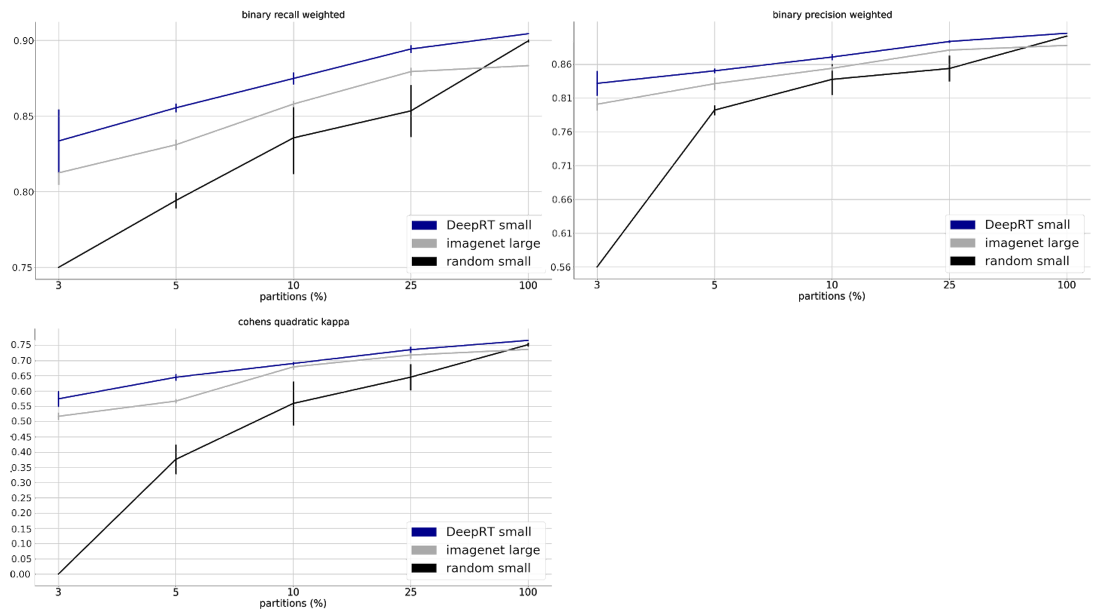


**Figure 8: Self supervised learning consistently outperforms random initializations and state of the art transfer learning across several outcome metrics:** *All outcome metrics analyzed for model performance. As in Fig.5, the x scale is displayed on logarithmic scale for visibility.*

The issue of overparameterization from state of the art models pre-trained on Imagenet becomes clear when observing train and validation cross entropy losses. The Imagenet ResNet50 model starts overfitting the data immediately as seen by the increased validation loss. At the same time the DeepRT model is able to continue to train and further minimize the validation loss (Fig. 9).

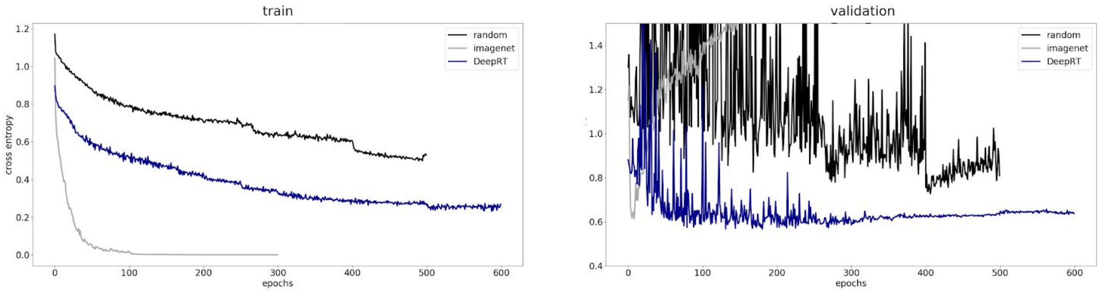


**Figure 9: Overparameterized imagenet model quickly overfits on smaller medical data sets:** *The train and validation loss shown over training epochs on the 10% partition. Here one sees immediate overfitting by the imagenet model.*

**Supplementary note 6: DeepRT not able to recover high resolution thickness maps in all cases.** While the average error was low for all test images, examples occured when DeepRT failed to accurately recover the retinal thickness maps directly from the fundus. In Figure 10 we see examples of top error and high error predictions. Notably in Fig. 10a the fundus images are visually unusual and the OCT derived thickness maps seem of lower quality. Also in Fig. b one of the fundus images is very dark. Despite the large data set used to train DeepRT, out of training distribution samples can occur and cause inaccurate predictions.

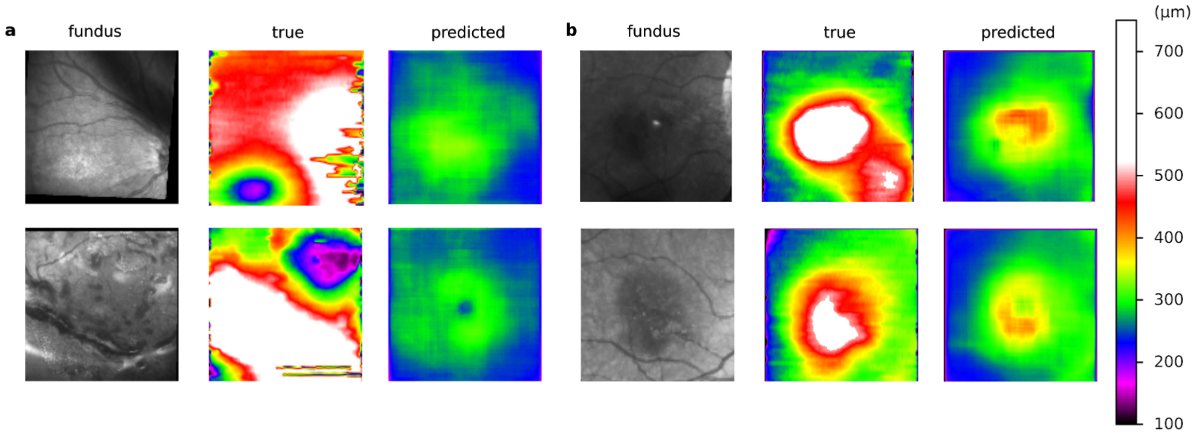


**Figure 10: High error predicted thickness maps:** *a. Top error thickness maps, MAE over 200 µm. b. High error examples, MAE over 80 µm*
